# A Change-point Analysis Method for Single-trial Study of Simultaneous EEG-fMRI of Auditory/Visual Oddball Task

**DOI:** 10.1101/100487

**Authors:** Aslan S Dizaji, Hamid Soltanian-Zadeh

## Abstract

Simultaneous EEG-fMRI has been vastly used to investigate functional networks of brain combining high spatial resolution (millimeter) of fMRI with high temporal resolution (millisecond) of EEG. However, to extract the most relevant information from the acquired data, it is necessary to develop analysis methods with less ad hoc assumptions. To this end, brain rhythms are often used which are the specific frequency-bands of EEG signal and assumed to represent diverse sub-second cognitive processes at different parts of the cortex. Furthermore, single-trial analysis of EEG is believed to show more realistic picture of ongoing and event-related activities of brain. Here, we present a nonparametric multiple change-point detection and estimation method for the single-trial analysis of simultaneous EEG-fMRI experiment recorded during auditory and visual oddball tasks. In a simple attention task like oddball, the frontal cortex of brain is responsible to distinguish and respond appropriately to target versus standard events. By using EEG signal at the frontal cortex, we show that the α-band activity changes according to “inhibition timing” hypothesis and the β-band activity is in line with “maintaining the status quo” hypothesis. Furthermore, using these activities to build regressors in the GLM analysis of fMRI, we localize active brain regions with high spatial and temporal resolutions and elaborate further on the coordination of attentional networks across brain.

## INTRODUCTION

Ongoing brain activity alternates in response to endogenous states and exogenous events. Electroencephalography (EEG) is one of the foremost tools to explore non-invasively this activity in human brain. Most notably, it is suitable for measuring the activity over the superficial layer of the scalp - cerebral cortex - with millisecond resolution. On the other hand, functional magnetic resonance imaging (fMRI) - another non-invasive neuroimaging technique - is able to localize active brain regions with unprecedented spatial resolution - millimeter - in the deep sub-cortical layers [1]–[3]. Simultaneous EEG-fMRI has been developed to utilize the complementary advantages of EEG and fMRI techniques and to decipher the coordination of functional networks across brain with high spatial and temporal resolutions [1] [3]. While the hardware design and data acquisition methods for these kinds of experiments have been already developed and in-use, there is not any universally standard approach for data analysis [4]–[6]. Some of these analysis methods use biophysical neurogenerative models to relate the hemodynamic events to EEG currents utilizing neurovascular coupling equations and parameters [3]. These models usually have many ad hoc assumptions and are computationally intensive. There are other kinds of methods, which use active brain regions obtained from fMRI as priors in the EEG source reconstruction [4]. These methods use parameterized forward head models and inverse modeling which are also computationally intensive. On the other hand, there are more data-driven approaches which combine symmetrically or asymmetrically EEG and fMRI data to find co-variation maps of them [1] [4]–[6]. One of such methods is EEG-informed fMRI regression which uses some features of multivariate EEG signal as regressors in the generalized linear model (GLM) to capture and isolate correlated patterns of fMRI signal of active brain regions [1] [4] [6].

The cerebral cortex plays a major role in all higher cognitive processes like attention, perception, decision-making, and planning. In a simple attention task like auditory/visual oddball, the most relevant part of brain which distinguishes target events from standard ones and plans desired actions, is frontal cortex. While traditional event-related potential can capture the average brain activity reliably, single-trial analysis of EEG signal recently has gained interest due to its ability to measure realistically the endogenous wax and wane of brain activity inherent to each trial [1]. Furthermore, it is believed that the frequency-band components of EEG signal have important implications on different cognitive processes. EEG is usually divided to five frequency-bands: δ-band (0.5–4 Hz), θ-band (4–8 Hz) α-band (8–14 Hz), β-band (14–30 Hz), and γ-band (30–100 Hz) [7]. Both amplitudes (powers) and phases of these frequency-band components carry information about cognitive processes inside brain [8]–[26]. As an instance, the α-band power, which is the most prominent EEG component particularly over the posterior part, is generally thought to correlate with inhibiting and idling state of the cortex and suppress task-irrelevant networks of brain [27] [28]. Based on “inhibition timing” hypothesis, the α-band activity represents suppression and selection, two important aspects of attention [29]. On the other hand, based on another hypothesis, the β-band activity, which its magnitude is more prominent in the sensorimotor cortex, is believed to signal “maintaining the status quo” in different parts of the cortex [30].

In this study, we develop a nonparametric method for single-trial analysis of EEG in the general framework of change-point analysis [31] [32]. While change-point analysis is a well-established method in signal processing; it has rarely used for EEG single-trial analysis. Moreover, our method is best suited for multidimensional data with unknown number and location of change-points by using minimum distribution assumptions. The estimation of change-points is based on U-statistics and hierarchical clustering with bisection approach for its computational efficiency [32]. We use this method to divide the multidimensional EEG signal of the electrodes of frontal cortex to distinct temporal windows and then calculate the single-trial relative powers of α and β-band components of these temporal windows. Our results provide support for α-band “inhibition timing” and β-band “maintaining the status quo” hypotheses. For single-trial analysis of simultaneous EEG-fMRI, we use α and β-band relative powers as regressors in the statistical parametric mapping of fMRI [33]–[35]. The results show the co-variation map of EEG and fMRI signals and they shed light on the spatial and temporal coordination of attentional networks in auditory and visual oddball tasks [36]–[38].

## MATERIALS AND METHODS

The EEG-fMRI data we use is from the simultaneous EEG-fMRI data collection described in [39]–[41], but we reproduce much of the relevant information here for ease of reading and refer the reader to those previous studies for further details. This data was obtained from the OpenfMRI database and its accession number is ds000116 (https://openfmri.org/dataset/ds000116).

### Auditory and Visual Oddball Paradigms

Seventeen subjects (six females; mean age of 27.7 years; age range of 20–40 years) participated in three runs of each of the auditory and visual oddball paradigms. The 375 (125 per run) total stimuli per task were presented for 200 ms each with a 2–3 s uniformly distributed variable inter-trial interval (ITI) and target probability of 0.2. The first two stimuli of each run were restricted to be standards. For the visual task, the target was a large red circle (3.45° visual angle) and the standard was a small green circle (1.15° visual angle), both of them on iso-luminant gray backgrounds. For the auditory task, the target sound was a broadband “laser gun” and the standard stimulus was a 390 Hz pure tone. Because our study is about task-related cognitive states, subjects were asked to respond to the target stimuli, using a button press with the right index finger on a button response pad.

### Simultaneous EEG-fMRI Data Acquisition

A 3 T Philips Achieva MRI scanner (Philips Medical Systems) was used to collect functional echo-planar image (EPI) data with 3 mm in-plane resolution and 4 mm slice thickness. It covered entire brain by obtaining 32 slices of 64*64 voxels using a 2000 ms repetition time (TR) and 25 ms echo time (TE). Also, high-resolution single-volumes of a 2*2*2 mm EPI image and a 1*1*1 mm spoiled gradient recalled image (SPGR) were obtained for each subject for the purpose of registration.

Simultaneously and continuously, EEG was recorded using a custom-built MR-compatible EEG system with differential amplifier and bipolar EEG cap. The caps were consisted of 36 Ag/AgCl electrodes including left and right mastoids, arranged as 43 bipolar pairs. Bipolar pair leads were twisted to minimize inductive pickup from the magnetic gradient pulses and subject head motion in the main magnetic field. This oversampling of the electrodes ensured that the data forms a complete set of electrodes even when there is a need to discard noisy channels. Due to the existence of magnetic field, the EEG signal is contaminated with gradient and Ballistocardiogram (BCG) artifacts. To enable removal of the gradient artifacts in our offline preprocessing, the 1-kHz-sampled EEG signal was synchronized with the MR-scanner clock at the start of each of 170 functional image acquisitions. A comprehensive description of the hardware and data acquisition can be found in [39]–[42].

### EEG Data Preprocessing

We followed closely the standard offline preprocessing of EEG described in [39]–[44] using MATLAB (MathWorks). First, we removed the gradient artifacts by subtracting the mean EEG signal across all functional volume acquisitions from the initial EEG signal. We then applied a 10 ms median filter to remove any residual spike artifacts [44]. Secondly, we used the following digital Butterworth filters in the form of a linear-phase finite impulse response (FIR): a 1 Hz high pass filter to remove direct current drift, 60 and 120 Hz notch filters to remove electrical line noise and its first harmonic, and a 100 Hz low pass filter to remove high-frequency artifacts not associated with neurophysiological processes. BCG artifacts share frequency content with EEG activity and existing BCG removal algorithms cause loss of signal power in the EEG. Therefore, we performed single-trial analysis on the event-related potential based on the change-point analysis method before BCG artifacts removal. However, to isolate the N100, P200, and P300 components (Fig. 1) and compute the scalp topographies (Figs. 2–5), BCG artifacts were removed from the EEG data using a principal components analysis method [42] [43]. First, the data were low pass filtered at 4 Hz to extract the signal within the frequency range in which BCG artifacts are observed and then the first two principal components were calculated. The projection of channel weightings corresponding to those components subtracted out from the broadband data. These BCG-free data were then re-referenced from the 43 bipolar channels to the 34 electrodes space to isolate the N100, P200, and P300 components (Fig. 1) and compute the scalp topographies (Figs. 2–5). By visual inspection, trials containing motion or blink artifacts and also those with incorrect responses, were discarded from both auditory and visual datasets.

**Fig. 1.**
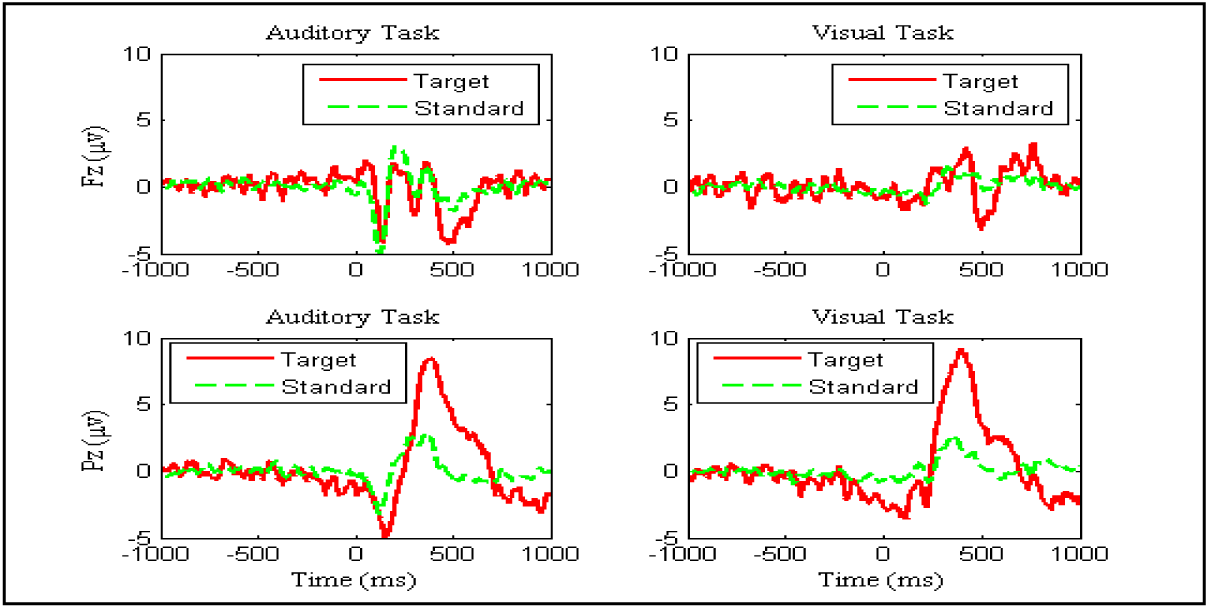
Traditional average event-related potential of Fz and Pz electrodes.

**Fig. 2.**
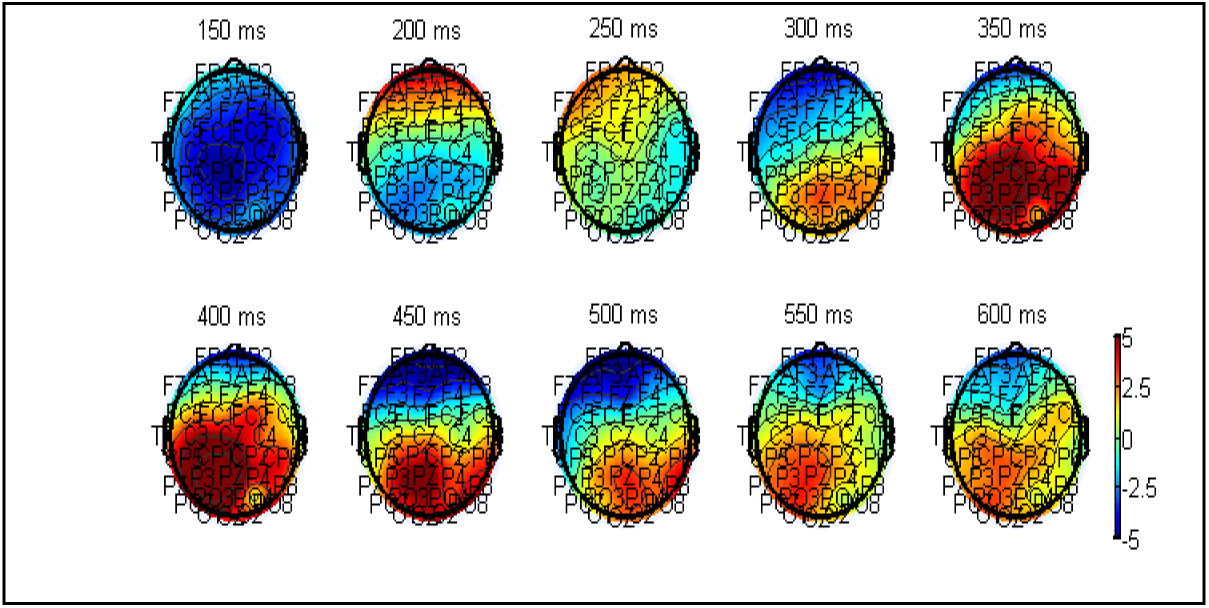
Average scalp topographies of auditory task for target trials.

**Fig. 3.**
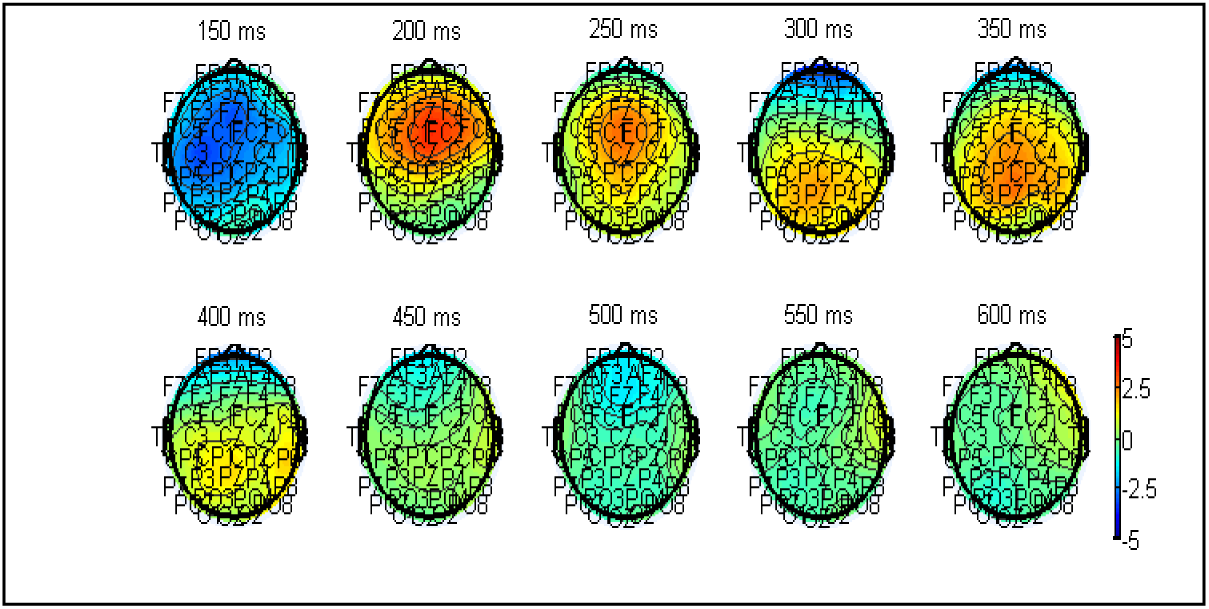
Average scalp topographies of auditory task for standard trials.

**Fig. 4.**
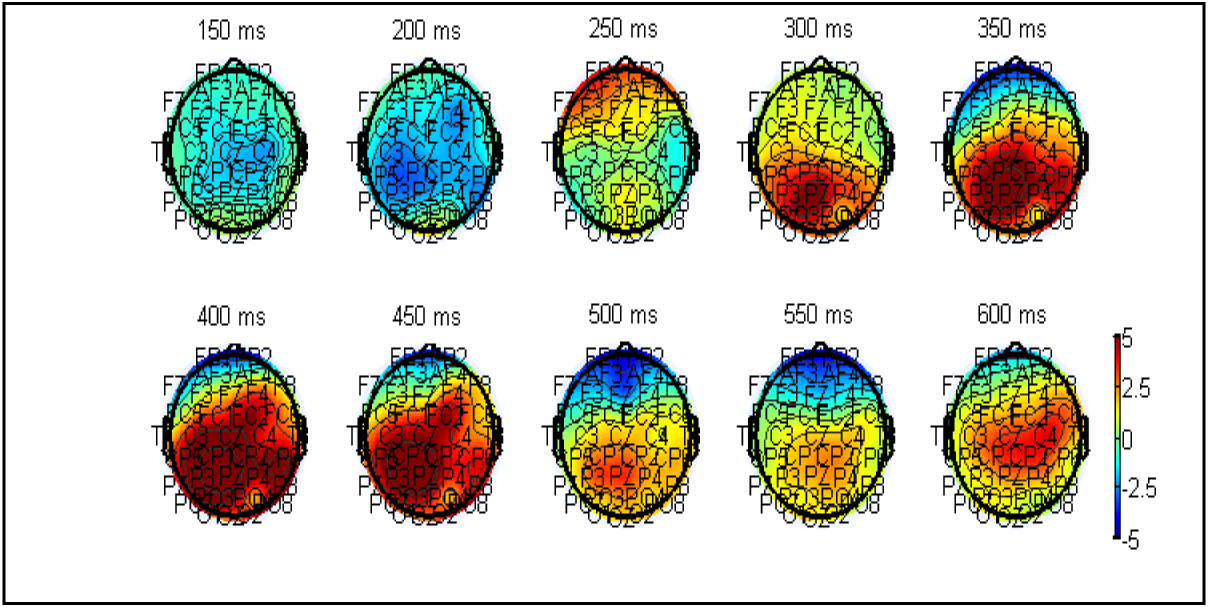
Average scalp topographies of visual task for target trials.

**Fig. 5.**
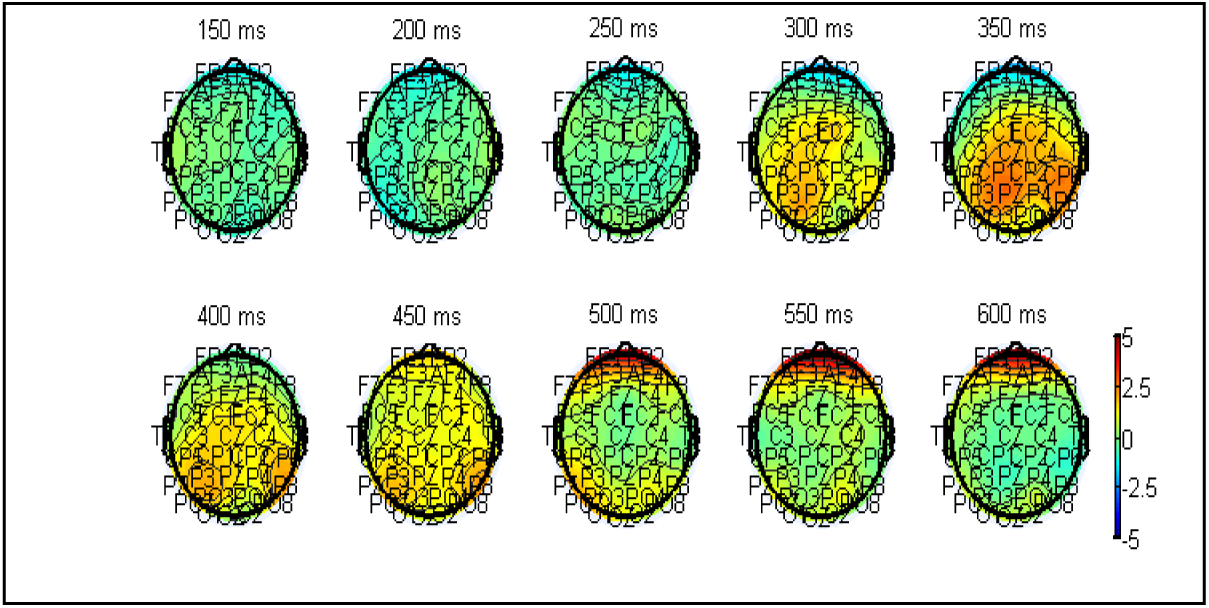
Average scalp topographies of visual task for standard trials.

### EEG Change-point Analysis

We defined the stimulus-locked −1000 ms to 1000 ms EEG epoch as a trial. We chose this interval because it ensures no overlap between adjacent trials due to ITI and also provides us with pre-stimulus potential for subsequent frequency-band analysis. We combined all target trials across three runs of each one of the auditory and visual tasks for each subject (at most 75 trials for each subject and task). We did the same for the standard trials (at most 300 trials for each subject and task).

We are interested in task-related EEG response which is the brain activity for distinguishing target from standard events and subsequent appropriate action. The EEG signal of the electrodes at the frontal cortex is believed to indicate this activity. As a result, we used the referencing matrix of each subject to transfer the EEG signal from the 43 bipolar channels to the 34 electrodes space and then chose 7 frontal cortex electrodes (Fp1, Fp2, AF3, AF4, F3, Fz, F4 in international 10–20 system) for change-point analysis.

We performed change-point analysis on average event-related potential obtained from 7 frontal cortex electrodes independently for each subject, task (auditory/visual), and trial type (target/standard). By doing this, we increased the signal-to-noise ratio of task-related EEG potential. However, our subsequent single-trial analysis based on frequency-band components was rested on individual trials. Also, to decrease the computational demand of our change-point analysis, we averaged the EEG signal over 10 ms sliding window, so instead of having EEG epochs with 2000-point length (1000 Hz), we had epochs with 200-point length (100 Hz) covering stimulus-locked −1000 ms to 1000 ms interval. We think this does not change the accuracy of our change-point analysis (particularly considering the low signal-to-noise ratio of the EEG datasets obtained in the MR environment), while it substantially increases the computational efficiency (Our change-point method runs at O(kT^2^); k is the number of the change-points and T is the sample-length of a signal).

Our change-point formulation is based on a recently developed nonparametric multiple change-point detection and estimation method which has just one assumption behind its applicability; the multidimensional signal must have μ-th absolute moment for some μ є (0,2) [32]. By combining it with hierarchical estimation (bisection method) and significance testing (permutation), we efficiently and reliably isolated up to 10 change-points from our EEG epochs.

In the following formula, μ = 1, T = 200 (the length of our EEG epoch), and Z_κ_ is a point in a 7-dimensional EEG epoch time-series in which κ є (1, 200). In (1), we vary κ along an EEG epoch and then move τ along the resultant temporal window, thus τ є (1, κ). As a result, we have two separate time-series with two probability distributions: F_x_ for X_τ_ and F_y_ for Y_τ_(κ). Based on this definition, we calculate *Ê* from (2) and *Q̂* from (3). According to (4), if τ is a change-point, then F_x_ is not equal to F_y_ and *Q̂* must go to infinity. Using (5), by spanning an EEG epoch with κ and τ, we are able to find the maximum possible value for *Q̂* and the corresponding estimated change-point τ. For more complete description, please refer to [32].

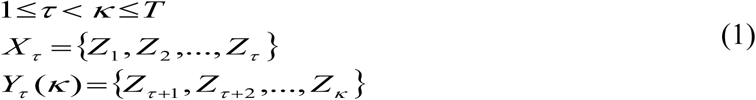

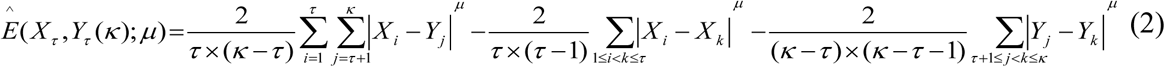

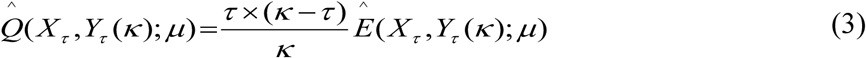

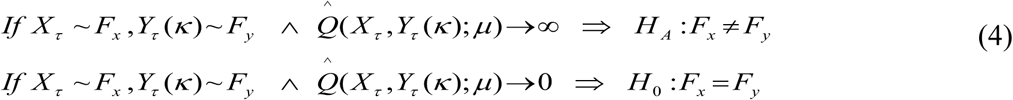

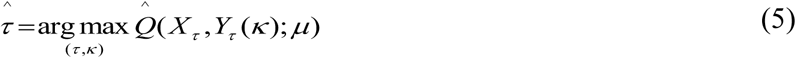

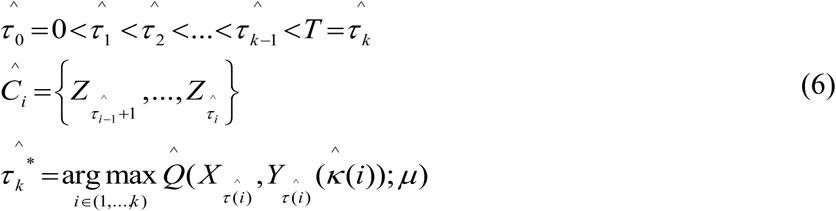

We performed this estimation hierarchically to find the desired number of significant change-points in our EEG epochs based on (6) (see Figs. 6–7 for average results across subjects). Also, to test the significance of our change-point detection, we performed permutation test. We permuted the signal values on our EEG epoch time-series (length = 200 points) 100 times for each subject, task, and trial type separately. Then, we ran the same hierarchical change-point detection and estimation procedure by using (6) for each permutation run (see Fig. 8 for average results across subjects). Finally, we compared the distribution *Q̂*_max_ of for permuted EEG epochs with the original EEG epochs to verify the significance of our detected change-points (see Table I for average results across subjects).

**Fig. 6.**
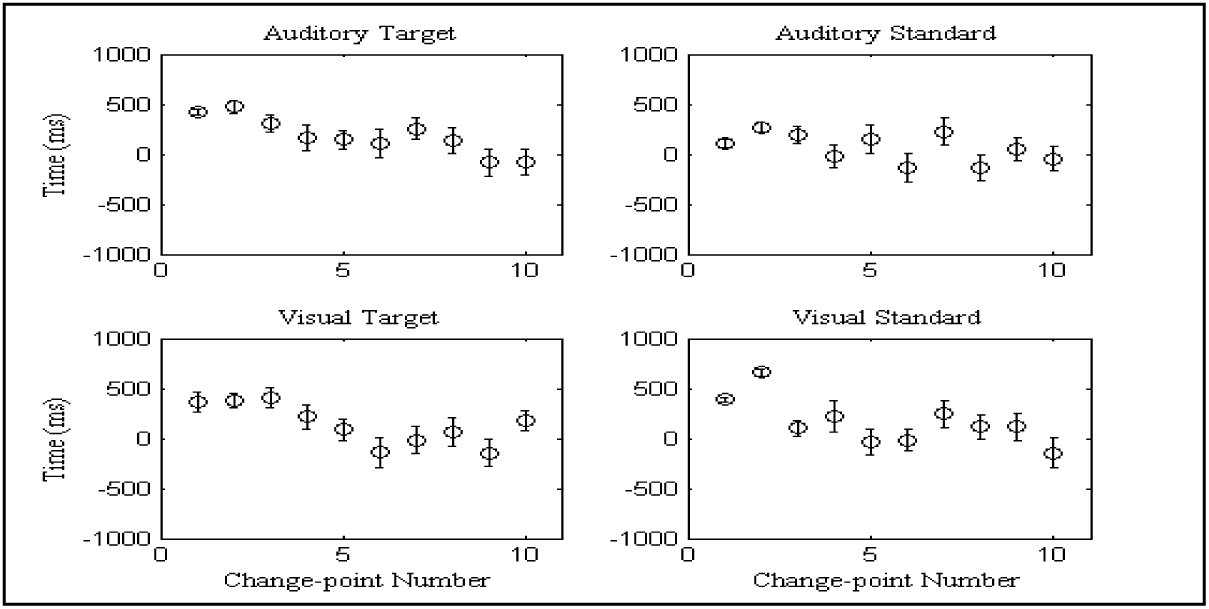
The first 10 change-points and their standard errors.

**Fig. 7.**
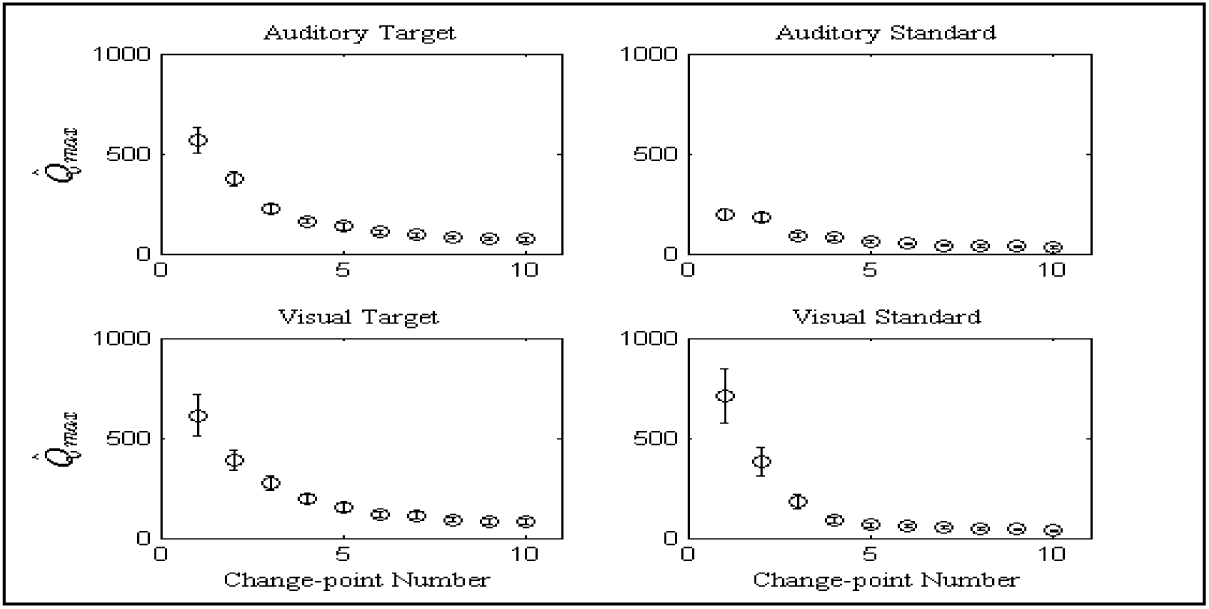
The *Q̂*_max_ values of the first 10 change-points and their standard errors.

**Fig. 8.**
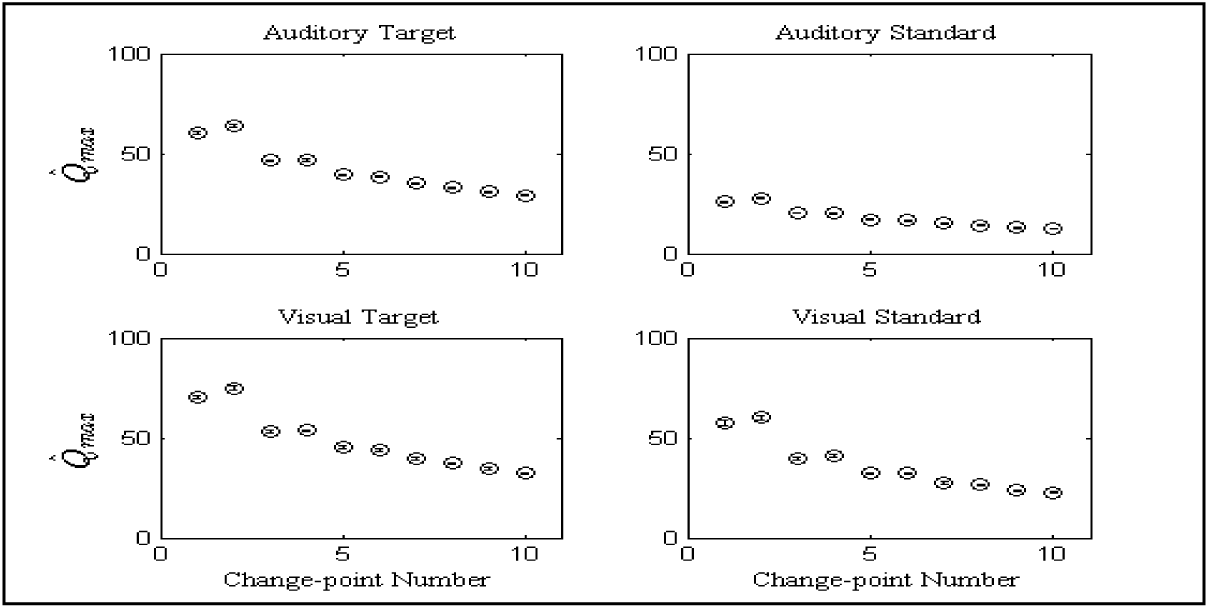
The *Q̂*_max_ values of the first 10 change-points of permuted trials and their standard errors.

**TABLE I:** The summary of change-point analysis averaged across all subjects for each combination of task and trial type.

| Change-points Number<br>(order of significance) | Auditory Target |  |  | Auditory Standard |  |  | Visual Target |  |  | Visual Standard |  |  |
| --- | --- | --- | --- | --- | --- | --- | --- | --- | --- | --- | --- | --- |
| | CP | $\hat{Q}_{\max}$ | $\hat{Q}_{\max}^p$ | CP | $\hat{Q}_{\max}$ | $\hat{Q}_{\max}^p$ | CP | $\hat{Q}_{\max}$ | $\hat{Q}_{\max}^p$ | CP | $\hat{Q}_{\max}$ | $\hat{Q}_{\max}^p$ |
| 1 | 426 | 566 | 60 | 105 | 197 | 26 | 363 | 613 | 70 | 387 | 712 | 57 |
| 2 | 474 | 374 | 64 | 267 | 179 | 28 | 380 | 392 | 75 | 659 | 382 | 60 |
| 3 | 309 | 223 | 47 | 194 | 93 | 20 | 409 | 275 | 53 | 104 | 181 | 40 |
| 4 | 161 | 163 | 47 | -18 | 79 | 20 | 219 | 193 | 54 | 219 | 91 | 41 |
| 5 | 148 | 136 | 40 | 146 | 60 | 17 | 87 | 157 | 45 | -34 | 66 | 32 |
| 6 | 107 | 109 | 38 | -131 | 51 | 16 | -142 | 117 | 44 | -15 | 59 | 32 |
| 7 | 256 | 95 | 35 | 229 | 41 | 15 | -16 | 114 | 40 | 244 | 56 | 27 |
| 8 | 136 | 80 | 33 | -135 | 39 | 14 | 66 | 93 | 37 | 116 | 49 | 27 |
| 9 | -83 | 73 | 31 | 51 | 36 | 13 | -145 | 82 | 35 | 117 | 43 | 24 |
| 10 | -79 | 73 | 29 | -43 | 32 | 12 | 180 | 80 | 32 | -145 | 37 | 23 |

### EEG (α, β)-band Single-trial Analysis

For single-trial analysis of EEG, we considered EEG α and β-band components due to their relevance to our simple paradigm and minimum contamination with BCG artifacts. After calculating the change-points of average event-related potential for each subject, task, and trial type separately; we returned to our target and standard single-trials and divided the stimulus-locked −1000 ms to1000 ms trial interval to distinct temporal windows each one of them starts at −1000 ms and ends at a change-point (CP). (We did this instead of considering the time-span between two consecutive change-points due to the fact that the change-points were close to each other and the time-gaps were not sufficient to let us reliably calculate the powers of α and β-band components.)

Afterward, we calculated average α and β-band powers for the temporal windows of each one of the trials, and the average broadband (BB) power (0–100 Hz) of the same trial. We divided the α and β-band powers of all temporal windows of a trial to the broadband power of the same trial by using (7) and obtained the relative powers of α and β-band components. By using this approach, we simultaneously achieved two objectives: 1) we normalized the α and β-band powers of the temporal windows of a trial on the same ground. 2) we removed any baseline effects of EEG signal so we were able to combine all the relative powers across all the trials for each subject, task, and trial type separately.

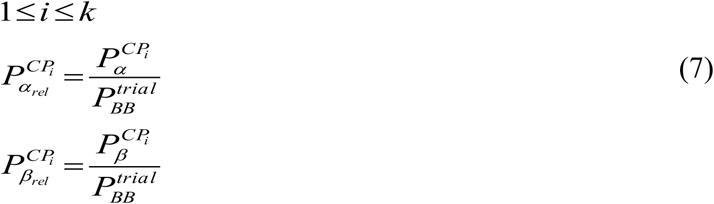

Furthermore, to have summary statistics of α and β-band relative powers across all temporal windows and subjects (but independently for each task and trial type), we divided the 2000-point length stimulus-locked EEG epoch to 20 equal bins each one has 100-point length. Then we placed the α and β-band relative powers of all subjects in one of those bins based on their temporal windows’ end positions. Afterward, for each bin, we calculated the mean and standard error of the mean for α and β-band relative powers. We presented these results in Figs. 9–12 independently for each task and trial type.

**Fig. 9.**
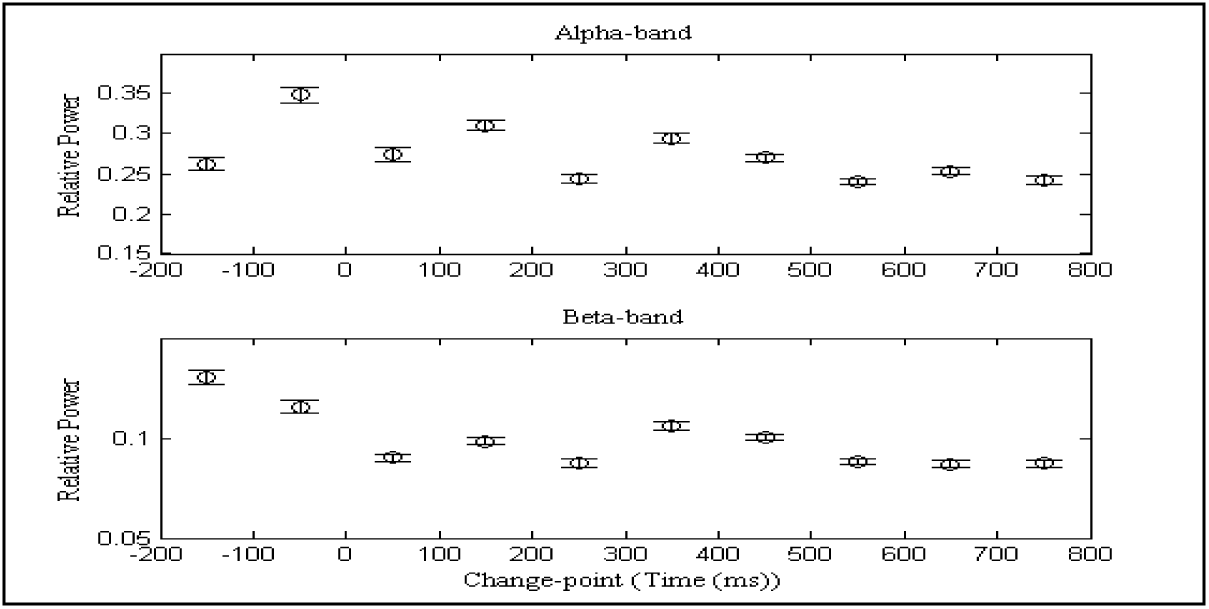
α and β-band relative powers of auditory task for target trials.

**Fig. 10.**
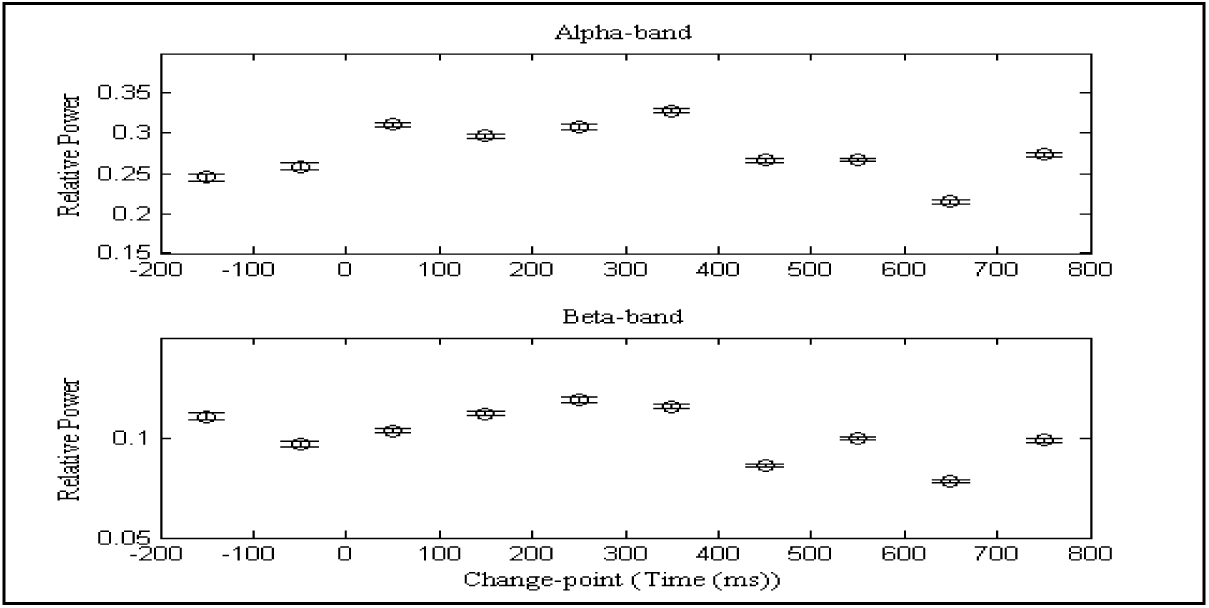
α and β-band relative powers of auditory task for standard trials.

**Fig. 11.**
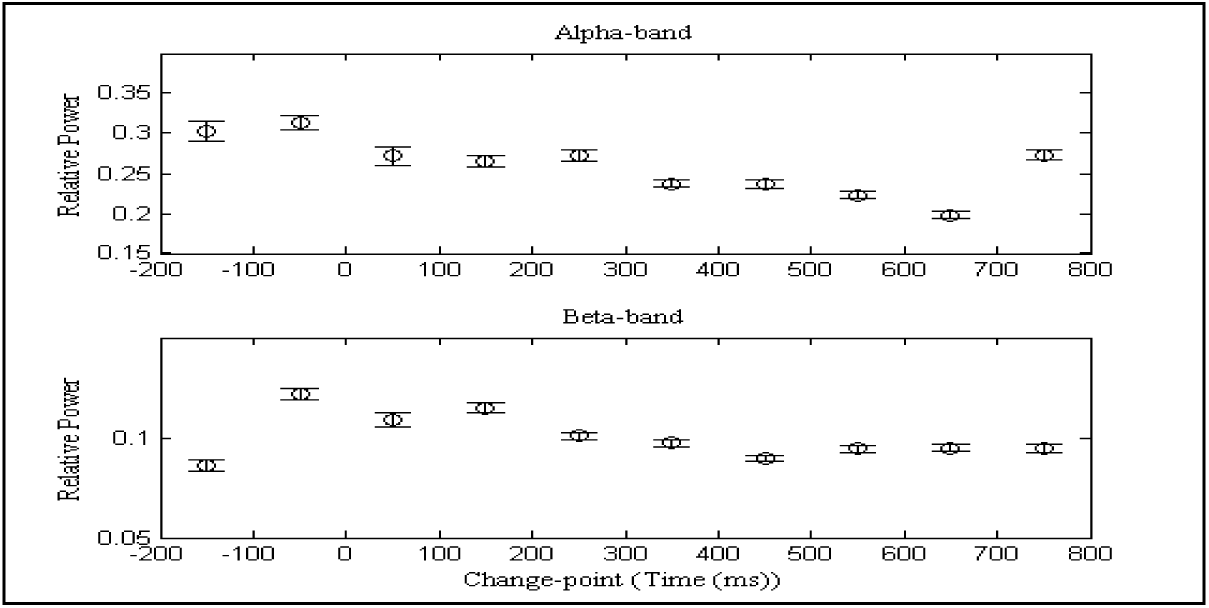
α and β-band relative powers of visual task for target trials.

**Fig. 12.**
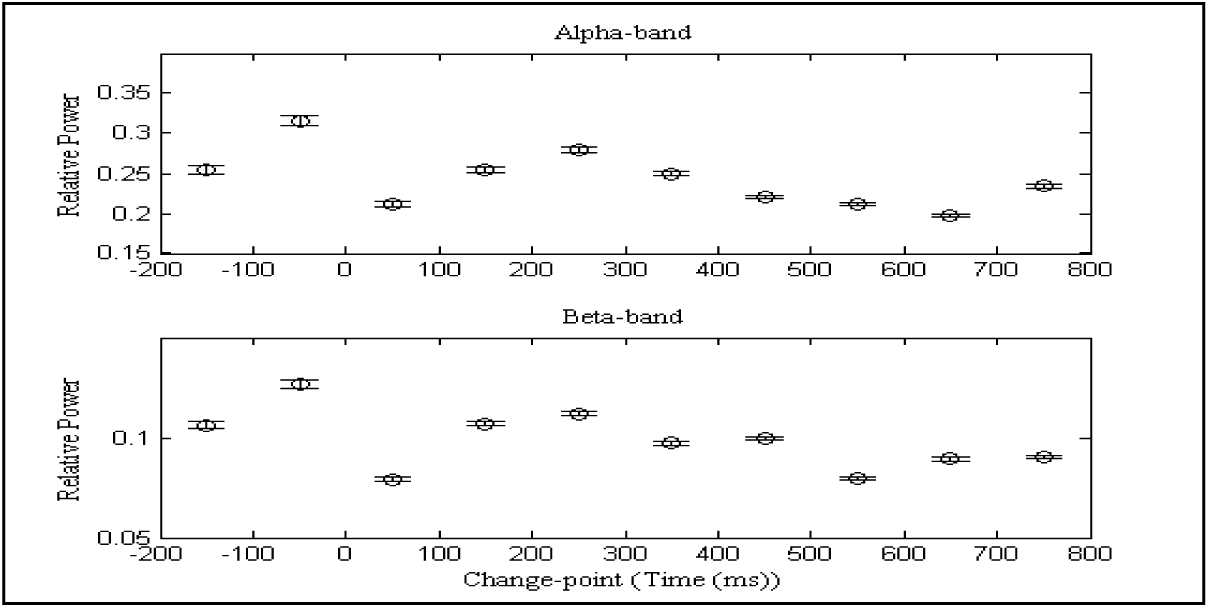
α and β-band relative powers of visual task for standard trials.

### fMRI Data Preprocessing

Using SPM8 (Wellcome Department of Imaging Neuroscience, London, UK; see http://www.fil.ion.ucl.ac.uk/spm), we performed realignment, un-warping, and slice-timing correction on the functional images for each subject, task, and run separately. We later included the realignment parameters as confounds of no interest in the subsequent GLM analysis of fMRI. After brain extraction, we co-registered the high resolution SPGR images (1*1*1 mm) to the high resolution EPI images (2*2*2 mm) and segmented the co-registered SPGR images to grey matter, white matter, and cerebrospinal fluid for each subject. Then, by using the resultant parameters, we normalized the realigned, un-warped, and slice-timing corrected functional images to the standard Montreal Neurological Institute (MNI) brain template for each subject, task, and run separately (we normalized the co-registered SPGR images to the standard MNI brain template for each subject too). Afterward, we spatially smoothed the functional images by using 8 mm full-width half-maximum (FWHM) Gaussian kernel for each subject, task, and run separately. Finally, for each subject, we checked manually the alignment of the smoothed functional images to the normalized SPGR images and we discarded the data of subject 13 due to misalignment.

### Traditional fMRI Analysis

In the traditional analysis of fMRI, we used multiple conditions files to feed target and standard event-related regressors to the SPM8 for GLM analysis. The event-related regressors comprised of boxcar functions with unit amplitude and onset and offset matching that of the stimuli with 200 ms duration. The regressors were already orthogonal to each other so we were able to capture the whole variability of each regressor completely [45]. Both regressors were convolved with the canonical hemodynamic response function and temporal derivatives were included as confounds of no interest (The other confounds of no interest were multiple regressors files of realignment parameters). Also, the functional data were high-pass filtered at 100 s and AR(1) was used to model serial correlations. Event-related target, standard, and target minus standard t-contrasts were constructed (replicated and scaled) and a fixed-effects model was used to model activations across runs for each subject and task separately. The one-sample t-test was used to compute the group-level mean activations of these contrasts across subjects for each task separately (the data of subject 13 was already removed). The t-maps of these group-level statistical images were thresholded at p-value = 0.01 and overlaid on the standard MNI brain template.

### EEG-based Single-trial Analysis of fMRI

For EEG-based single-trial analysis of fMRI, we used multiple conditions files to feed the α and β-band relative powers of all temporal windows of each subject and task as parametric modulators in the SPM8 for GLM analysis. So for each subject and task, we have 20 additional regressors based on 2 frequency-bands and 10 change-points. These EEG-based regressors were designed with 100 ms duration centered on the change-points and their heights were modulated using the α and β-band relative powers of each temporal window of each trial and they were orthogonalized with respect to the traditional regressors by SPM8 internal capability. These regressors were convolved with the HRF and temporal derivatives were included as confounds of no interest (The other confounds of no interest were multiple regressors files of realignment parameters). Also, the functional data were high-pass filtered at 100 s and AR(1) was used to model serial correlations. For each subject, task, frequency-band, and temporal window, the following t-contrasts were constructed: single-trial target, single-trial standard, single-trial target minus traditional target, and single-trial standard minus traditional standard. These contrasts were replicated and scaled and a fixed-effects model was used to model the activations across runs for each subject, task, frequency-band, and temporal window separately. To have summary statistics of the single-trial effects of α and β-band relative powers of the EEG signal on the fMRI data across all temporal windows and subjects (but independently for each task), we divided the 2000-point length stimulus-locked EEG epoch to 20 equal bins each one has 100-point length and placed the t-contrasts of all subjects in one of those bins based on their temporal windows’ end positions. The one-sample t-test was used to compute the group-level mean activations of these contrasts for each task, frequency-band, and bin separately across subjects (the data of subject 13 was already removed). The t-maps of these group-level statistical images were thresholded at p-value = 0.01 and overlaid on the standard MNI brain template.

## RESULTS AND DISCUSSION

All subjects responded with high accuracy and speed. For the auditory task, 98.3 ± 2.0 *%* of targets was correctly detected with 404.1 ± 58.3 ms reaction time (RT) and for the visual task 98.4 ± 3.1 % of targets was correctly detected with 397.2 ± 38.9 ms RT.

### Traditional Event-related Potential

The average (across all subjects) event-related potential (ERP) which spans stimulus-locked −1000 ms to 1000 ms interval for Fz and Pz electrodes were displayed in Fig. 1 (independently for each task and trial type). P300 component which is an endogenous potential elicited in the process of attention, categorization, and decision-making is more prominent in the parietal sites between 300 ms to 500 ms. On the other hand, N100 and P200 are more prominent in the frontal sites. They are all visible for average ERP of target trials for both tasks in Fig. 1.

Also, the scalp topographies of average ERP across all subjects were shown in Figs. 2–5 in 50 ms intervals. In target trials with the right index finger response, we observe contralateral activity in the left motor cortex at the posterior parietal sites (Figs. 2 and 4). Also, for the auditory task, we have more activation in the right and left temporal sites due to the activity of auditory cortex (Figs 2–3) and for the visual task we see more activity in the occipital sites where visual cortex is engaged (Figs. 4–5).

### EEG Change-point Analysis

The average locations (across all subjects) of the first 10 change-points in −1000 ms to 1000 ms stimulus-locked interval were displayed in Fig. 6 (independently for each task and trial type). The first change-point of target trials for both tasks is approximately matched with the behavioral response or RT. In Fig. 7, the *Q̂*_max_ values of the detected change-points were plotted. The plot has a plain trend starting from large values and decreasing gradually. Also, in Fig. 8, the *Q̂*_max_ values of the first 10 change-points of permuted trials were plotted and, as expected, they have significantly lower *Q̂*_max_ values compare to Fig. 7.

We brought the summary of change-point analysis averaged across all subjects in Table I. For each combination of task and trial type, we have three columns: CP of stimulus-locked epochs in millisecond, *Q̂*_max_ values for the detected change-points, and *Q̂*_max_^*p*^ values for the permuted trials. The results clearly show that the detected change-points are significant for both tasks (auditory/visual) and both trial types (target/standard).

### EEG (α, β)-band Single-trial Analysis

We summarized the average results across all temporal windows and all subjects (the data of subject 13 was already removed) but independently for each task, trial type, and frequency-band component in Figs. 9–12.

In Figs. 9 and 11, for auditory and visual target trials respectively, the α-band activity is higher at the beginning and gradually decreases as the change-point moves to the right in the stimulus-locked EEG epoch. This is in accordance with the “inhibition timing” hypothesis [29] which states that the α-band power is negatively correlated with brain activity; it is higher at the pre-stimulus interval but it decreases as the frontal cortex gets engaged with task-related decision-making regarding the response to the target trials. The β-band activity is also in agreement with the “maintaining the status quo” hypothesis [30] which states that the β-band power is lower when we expect some cognitive processes happen which change the status quo. Here, in response to the target trials, the frontal cortex, which is the center of executive functions in brain, gets active to coordinate the proper action.

On the other hand, for auditory and visual standard trials in Figs. 10 and 12 respectively, we observe the opposite trend. Although, they start with relatively high pre-stimulus α and β-band activity, they decrease and again increase around 200 ms to 400 ms post-stimulus which is the time when brain is already processed the stimulus and the appropriate decision is made. This is again in line with the “inhibition timing” hypothesis for the α-band and the “maintaining the status quo” hypothesis for the β-band; in the standard trials, the frontal cortex is not involved in the coordination of any action and it just needs to keep the status quo which is the situation without any behavioral response.

To summarize this section, our developed change-point method and single-trial analysis are able to provide additional evidences for two relevant hypotheses in the literature related to α and β-band activities; they show how the brain rhythms in the task-related region of the cortex coordinate temporally with sub-second resolution.

### EEG-based Single-trial Analysis of fMRI

In this section, we brought a few plots from our single-trial EEG-fMRI analysis using α and β-band relative powers as regressors in the GLM and further elaborate on the relevancy of our results.

Fig. 13 shows the negative correlation of α-band activity of the electrodes of frontal cortex at the 50 ms temporal window with the BOLD response in auditory cortex in target trials. Also, the α-band activity of the same electrodes at the same temporal window has positive correlation with the BOLD response in thalamus and negative correlation with the BOLD response in brainstem in standard trials. These results corroborate the role of thalamo-cortical loop in attentional modulation and generation of α-band oscillations in cortical regions [20] [24] and “inhibition timing” hypothesis [29].

**Fig. 13.**
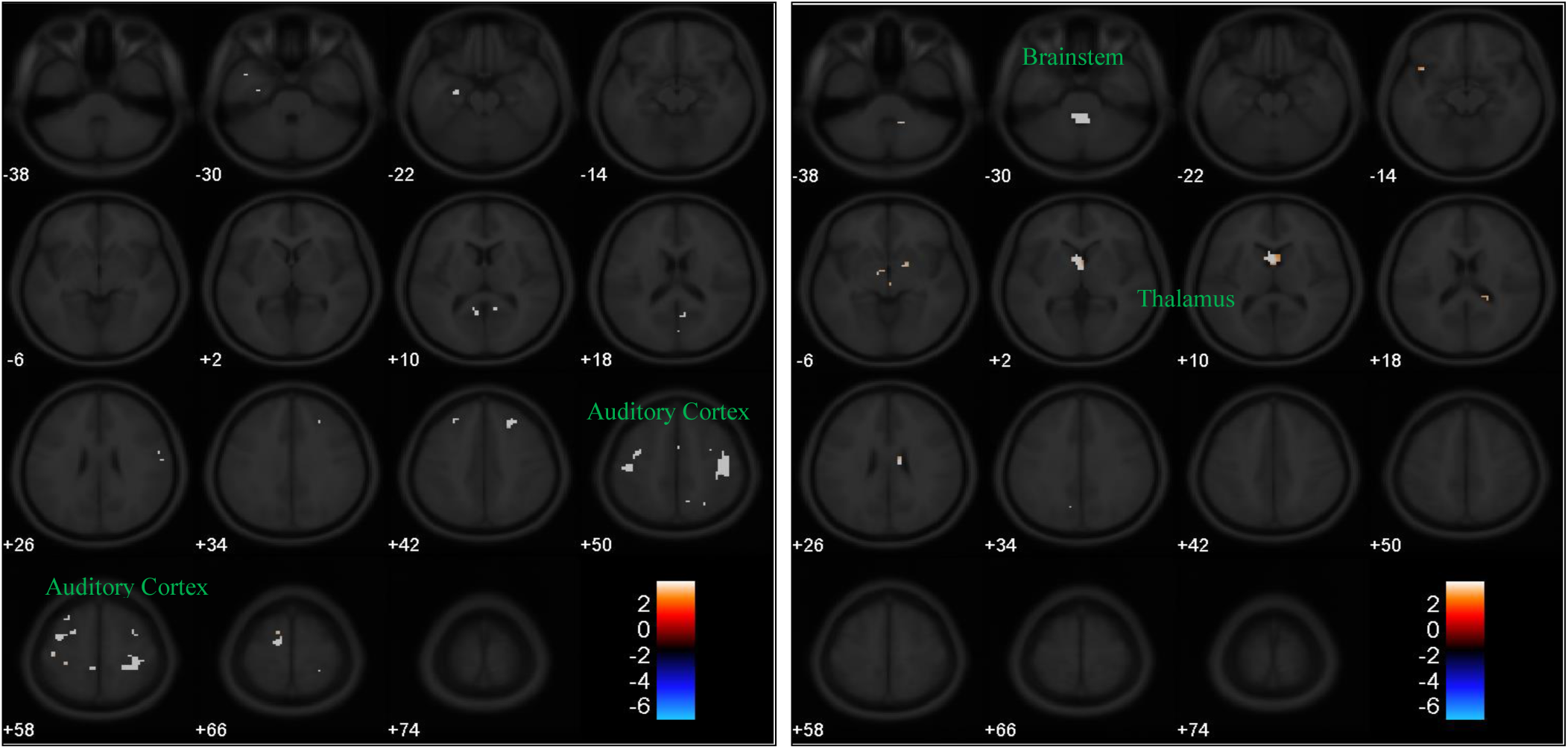
Pure effect of α-band relative power of 50 ms temporal window on BOLD responses of auditory target (left) and standard (right) trials.

Figs. 14 and 15 again show the positive correlation of α-band activity of the electrodes in frontal cortex with the BOLD response in thalamus in standard trials at the 150 ms and 450 ms temporal windows respectively. Also, there is a positive correlation between the α-band activity of the electrodes in frontal cortex with the BOLD response in cingulate cortex in standard trials at the 450 ms temporal window. Cingulate cortex is a part of the default mode network and this positive correlation shows the introspective evaluation of decisions in that region [40]. As it is clear from Figs. 13, 14, and 15, the α-band activity of the electrodes in frontal cortex has positive correlation with the BOLD response in thalamus and negative correlation with the BOLD response in brainstem only in standard trials. We believe this is due to the fact that there is higher inhibition in cortical regions during standard trials due to the lack of behavioral responses [39]–[41].

**Fig. 14.**
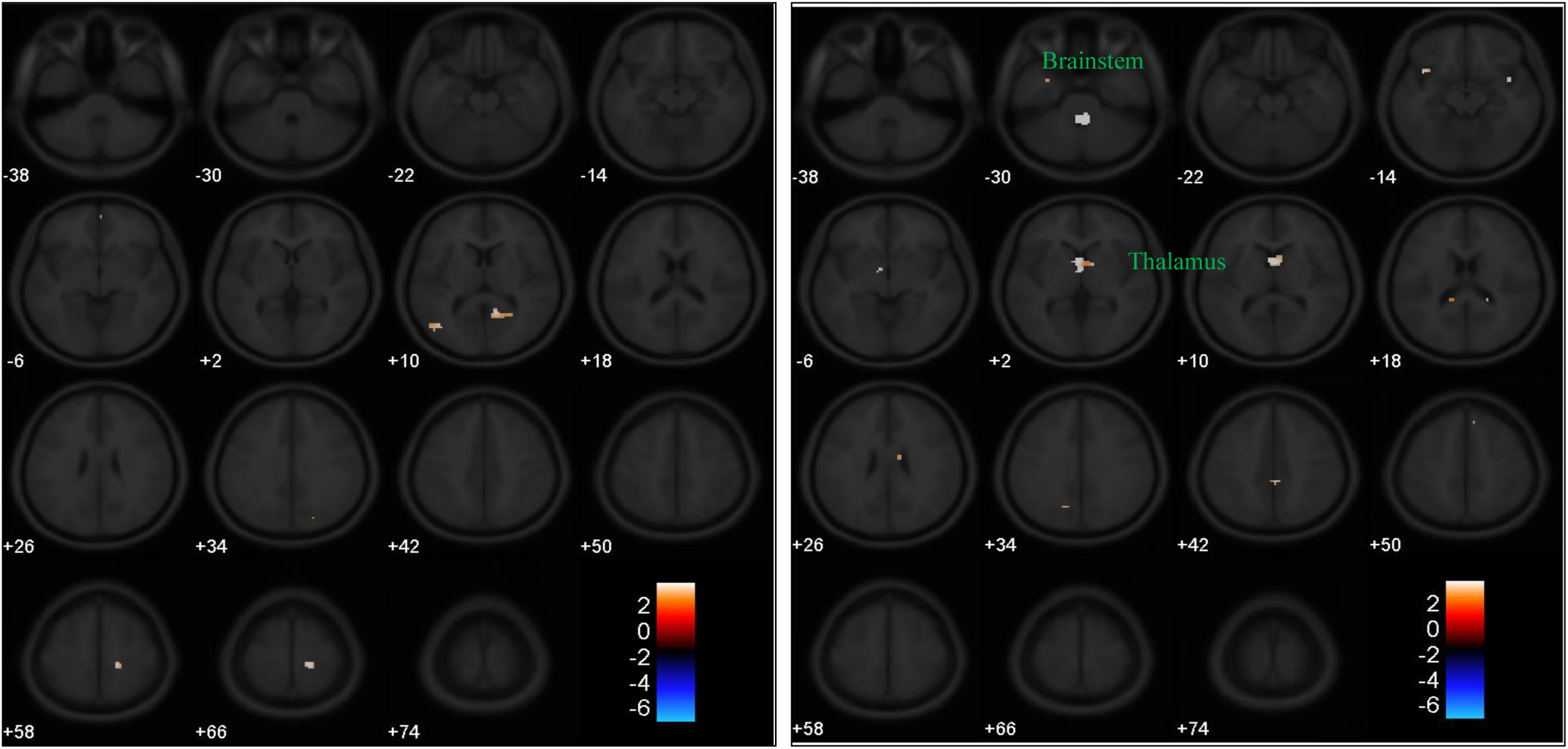
Pure effect of α-band relative power of 150 ms temporal window on BOLD responses of auditory target (left) and standard (right) trials.

**Fig. 15.**
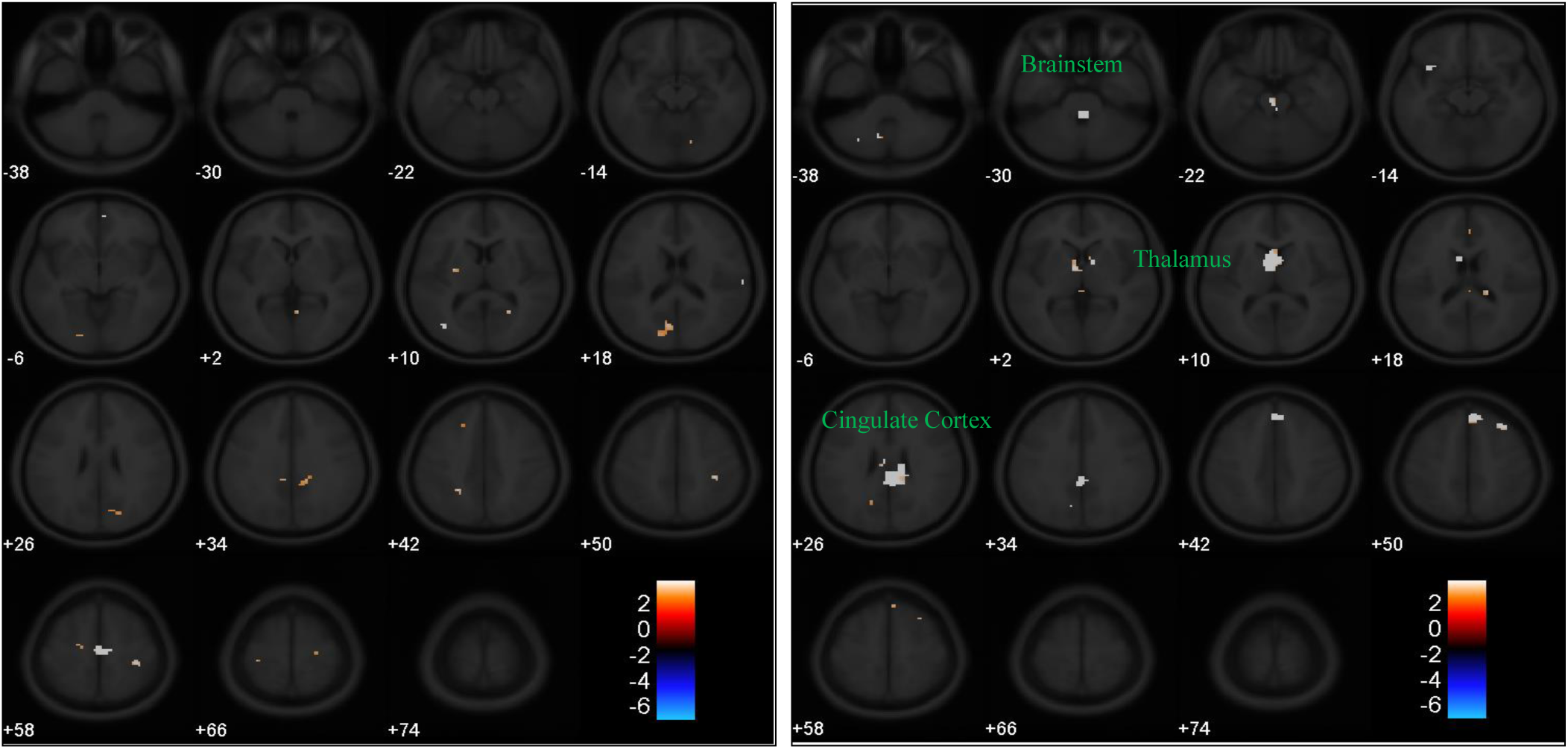
Pure effect of α-band relative power of 450 ms temporal window on BOLD responses of auditory target (left) and standard (right) trials.

Fig 16. shows the positive correlation of β-band activity of the electrodes in frontal cortex with the BOLD response in prefrontal areas in standard trials at the 150 ms temporal window. This further corroborates the role of β-band activity in temporal and spatial dynamics of the accumulation and processing of evidence leading to decision [8].

**Fig. 16.**
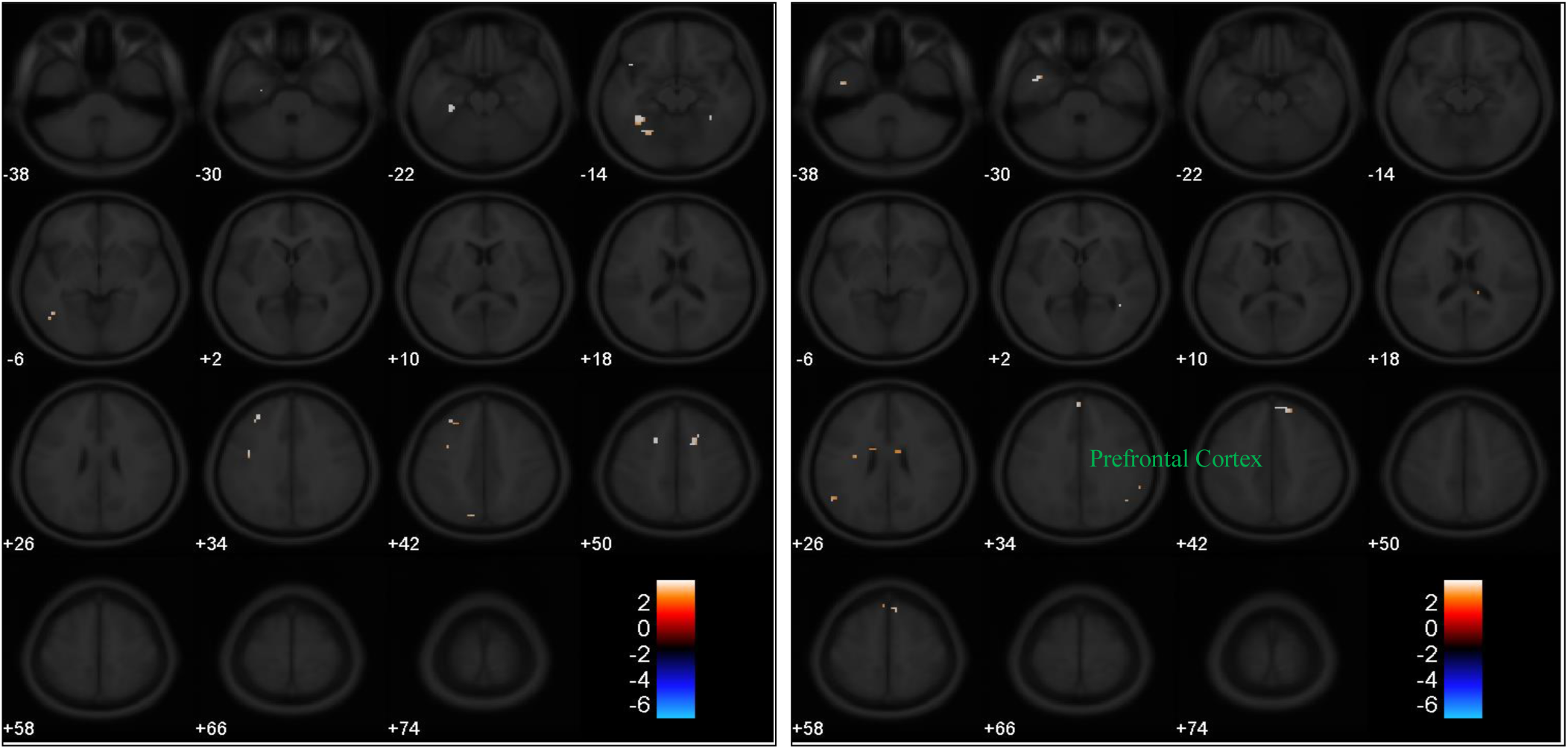
Pure effect of β-band relative power of 150 ms temporal window on BOLD responses of auditory target (left) and standard (right) trials.

Finally, Figs. 17 and 18 show the positive correlation of α-band activity of the electrodes in frontal cortex with the BOLD response in precuneus, a part of the default mode network, in target trials at the 350 ms and 550 ms temporal windows respectively. This positive correlation could be interpreted as the activation of the default mode network during and after behavioral responses and getting involved in the reflective self-awareness and introspective evaluation which are functions of precuneus [39]–[41].

**Fig. 17.**
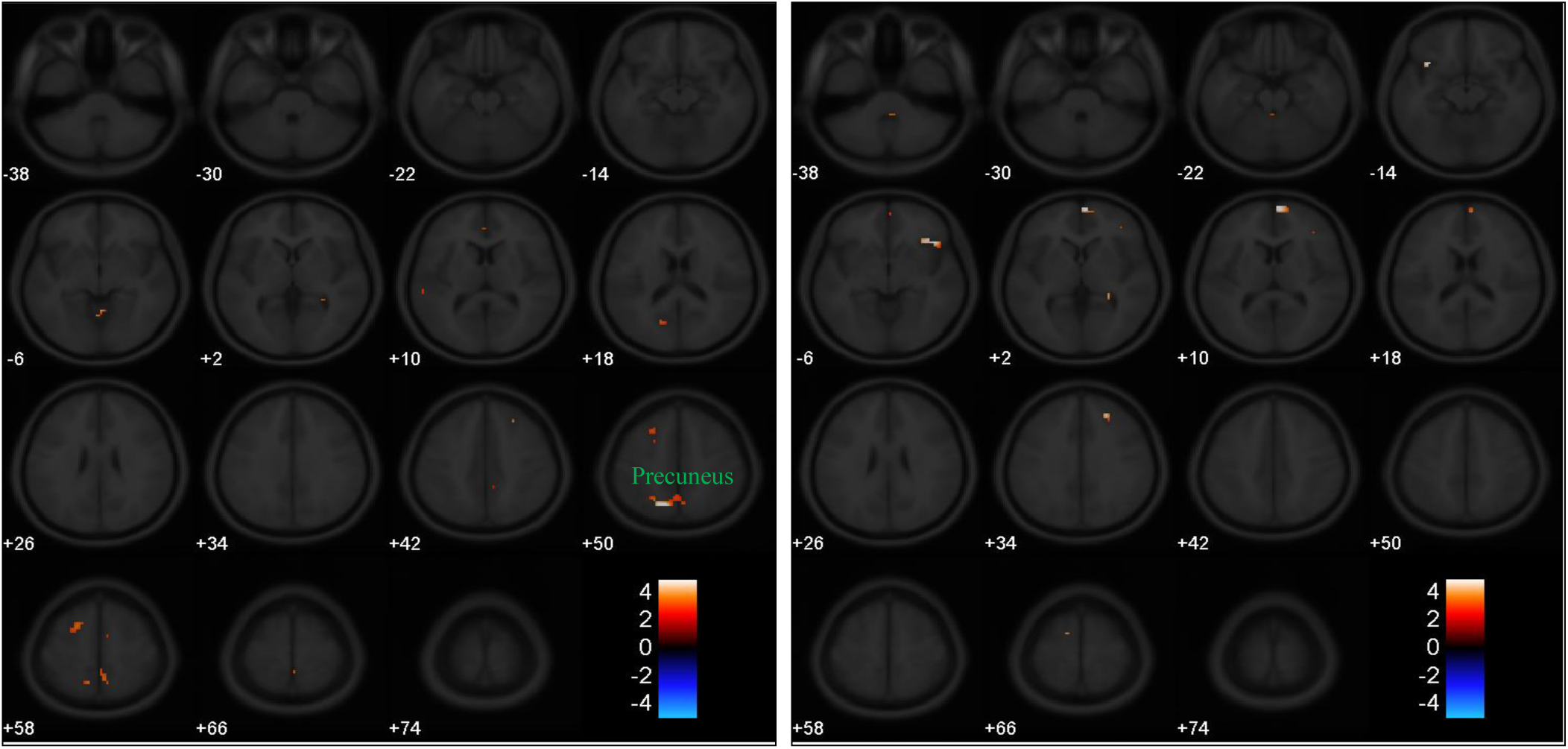
Pure effect of α-band relative power of 350 ms temporal window on BOLD responses of visual target (left) and standard (right) trials.

**Fig. 18.**
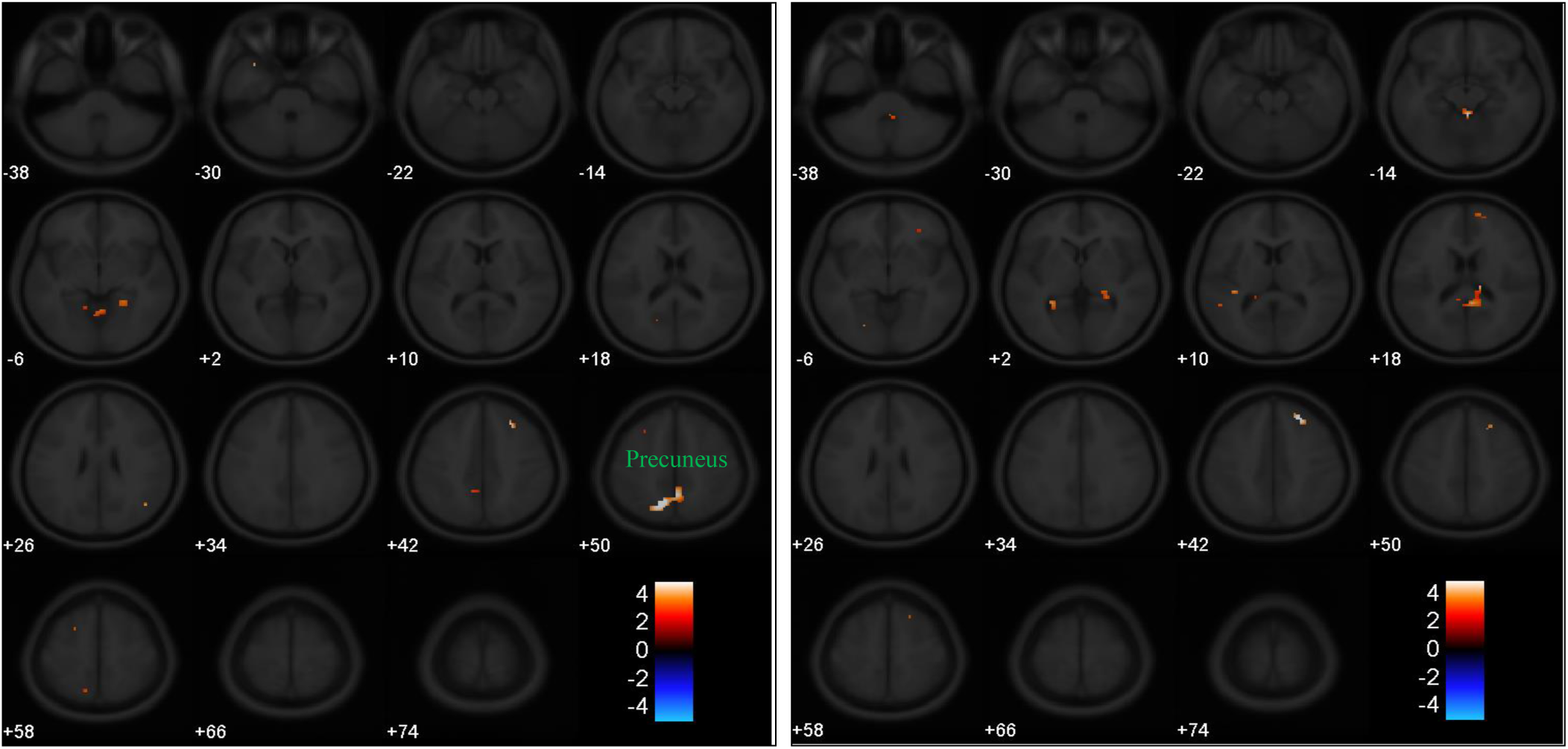
Pure effect of α-band relative power of 550 ms temporal window on BOLD responses of visual target (left) and standard (right) trials.

## ACKNOWLEDGMENT

A. S. D. was partly funded by the Iran’s National Elites Foundation. Also, we would like to thank HaDi MaBouDi and Safura Rashid Shomali for fruitful discussions.

